# Swim bladder resonance enhances hearing in crucian carp (*Carassius auratus*)

**DOI:** 10.1101/2022.08.01.502303

**Authors:** Hongquan Li, Zhanyuan Gao, Zhongchang Song, Yingnan Su, Wenzhan Ou, Jinhu Zhang, Yu Zhang

## Abstract

Sound sensing is vital for fish and more effort is necessary to address the hearing mechanism in fish. Here, we performed auditory evoked potentials (AEP) measurement, micro-computed tomography (Micro-CT) scanning, and numerical simulation to investigate the resonance of swim bladder and its influence on auditory sensitivity in crucian carp (*Carassius auratus*). The AEP results showed that at the tested frequency range up to 1000 Hz, the mean auditory thresholds of control fishes with an intact swim bladder were lower than that of treated fishes with a deflated swim bladder by 0.38–30.52 dB re 1 μPa. At the high frequency end, control fishes had a high but measurable auditory threshold. Correspondingly, numerical simulations showed that the intact swim bladder had a mean resonance frequency of 826±13.6 Hz, ranging from 810 to 840 Hz while the deflated swim bladder had no predominant resonance peak below 1000 Hz. The amplitude of received sound pressure at the resonance frequency for a sample in control group was 34.3 dB re 1 μPa higher than that for a treated sample, and the acceleration at the asteriscus of the control fish was higher than the treat fish by 43.13 dB re 1 m s^-2^. Both AEP experiment and modeling results showed that hearing sensitivity is enhanced through resonance of swim bladder in crucian carp and provided additional understandings on hearing mechanism in fish.

**Summary statement:** We used AEP measurement, Micro-CT scanning, and numerical simulation to demonstrate that the resonance of swim bladder can enhance hearing in crucian carp.

## INTRODUCTION

The ocean is a rich natural reservoir of sounds from biological, geophysical, and anthropogenic sources (Krause, 2008). The diversity of sounds provides ocean inhabitants with information and cues that facilitate survival. Many taxa, including fish, marine mammals, and invertebrates, have long been known as experts in producing and interpreting sounds to fulfill the biological processes of feeding, spawning, and social communication (Giorli et al., 2016; Hawkins and Amorim, 2000; Mann and Lobel, 1997; van Oosterom et al., 2016). The pressure and particle motion components of sound can be informative to animals that can sense sounds (Sueur and Farina, 2015; Thode et al., 2019). Sound sensing has been attributed to the development of hearing organs, such as statocysts in squids, and three pairs of otoliths and the lateral line in fishes (Montgomery et al., 2006; Mooney et al., 2010). These structures are key to collecting acoustic cues in animals that are sensitive to sounds. Among such animals, fish has long been known for their ability to detect two components of sound, namely scalar pressure and the back-and-forth vibration known as particle motion (Montgomery et al., 2006; Rogers and Cox, 1988).

Sound detection is critical for fish to settle in suitable habitats (Gordon et al., 2018; Parmentier et al., 2015), communicate (Maiditsch and Ladich, 2022), and avoid predators (Blaxter and Fuiman, 1990; Miller et al., 2006). The inner ear of fish is a multisensory organ consisting of three semicircular canals and three otolith end organs (Popper and Fay, 1993; Popper and Fay, 1997; Schulz-Mirbach et al., 2019). For teleosts, sound stimuli can reach the inner ear through direct and indirect pathways (Schulz-Mirbach et al., 2019). The otolith end organs (the utricle, saccule, and lagena) serve to detect linear acceleration in the direct pathway (Fig. 1A; Fay, 1984; Schulz-Mirbach et al., 2019; Sisneros and Rogers, 2016; Vetter and Sisneros, 2020). The calcareous otoliths with a higher density respond slower response to sound stimuli than tissue, resulting in a vibration difference that bends sensory hair cells to realize sensing (Hudspeth, 1985). All fish are thought to be able to detect particle motion through the direct pathway. Besides, most teleost species, such as goldfish (*Carassius auratus*) and walking catfish (*Clarias batrachus*), have evolved to own a swim bladder or other gas-filled cavities to detect sounds through the indirect pathway (Fig. 1B; Schulz-Mirbach et al., 2019; Shao et al., 2014). In this pathway, air cavities can oscillate to transduce sound energy to the inner ear (Popper et al., 2003; Schulz-Mirbach et al., 2019). Fish that can hear *via* both pathways (Fig. 1C) tend to have a wider auditory bandwidth and a higher sensitivity to sounds (Ladich, 2016; Popper and Fay, 2011).

**Fig. 1.**
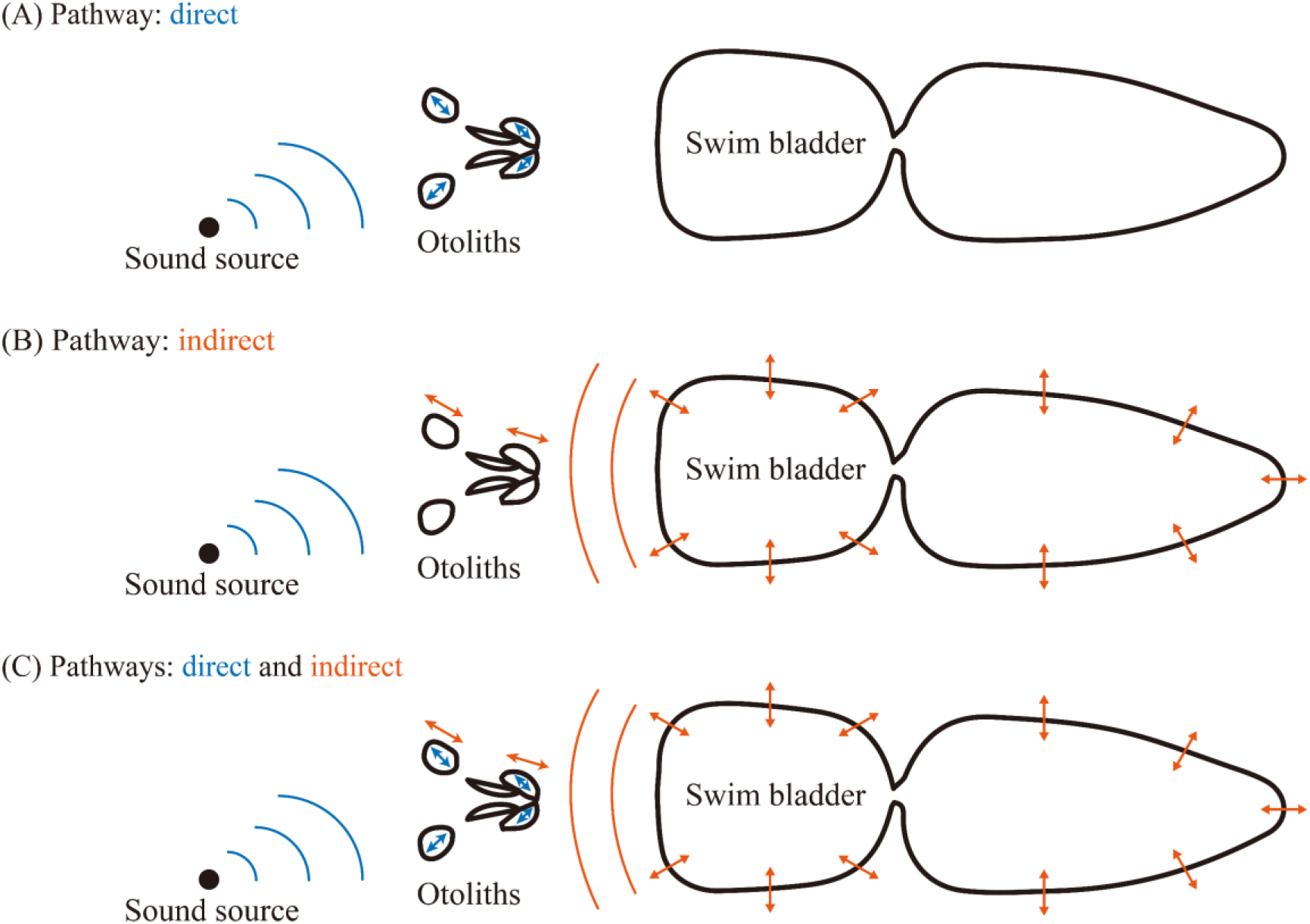
Modes of sound reception pathways in teleost fish with a swim bladder. (A) Stimulation of the otoliths through the direct pathway and (B) indirect pathway in which the swim bladder induces sound energy to the otoliths. (C) Sound sensing through the direct and indirect pathways.

Hearing capability can be examined through auditory evoked potentials (AEP) measurement (Corwin et al., 1982; Horodysky et al., 2008; Kenyon et al., 1998). Using this non-invasive electrophysiological method, the auditory evoked potentials from eighth nerves can be recorded to examine hearing in fish (Ladich and Fay, 2013) and investigate the role of auditory specializations (e.g., swim bladder) in hearing (Shao et al., 2014; Yan et al., 2000). The otolith end organs act as biological accelerometers that allow fish to directly detect particle motion (Vetter and Sisneros, 2020). Swim bladder functions as a pressure-to-particle motion transducer that lowers the hearing threshold and expands the auditory bandwidth (Schulz-Mirbach et al., 2019). The morphological features of the swim bladder and their contribution to hearing have been investigated in previous studies (Colleye et al., 2019; Schulz-Mirbach et al., 2012; Vetter and Sisneros, 2020; Yan et al., 2000), suggesting the removal of swim bladder resulted in high hearing threshold in plainfin midshipman (*Porichthys notatus*). But in oyster toadfish (*Opsanus tau*), the deflation of swim bladder did not influence hearing sensitivity (Yan et al., 2000). The variability across species requires more research to address the role of swim bladder in fish hearing and the potential mechanism hidden behind (Salas et al., 2019).

To gain more insights into the hearing mechanism in fish, we characterized the audiograms of crucian carps with an intact swim bladder (control group) and those with a deflated swim bladder (treated group) by AEP measurement. Micro-CT scanning, reconstruction, and numerical simulations were presented to provide additional data to elucidate the influence of swim bladder on hearing and sound reception in crucian carp.

## MATERIALS AND METHODS

### Sample preparation

Ten crucian carp samples were purchased from a local market and maintained in a rectangular freshwater tank with a high dissolved oxygen concentration for at least three days before experiment. The temperature was tracked daily, and all samples were exposed to natural light. Prior to the experiment, fork length (FL) was measured. The samples used in subsequent AEP measurements were divided into the control group and treated group: five samples with an intact swim bladder (control group) and five samples with a deflated swim bladder (treated group).

Before each AEP trial, the treated fishes were anesthetized with a solution of 100% clove oil (0.05–0.07 ml l^-1^, dependent on sample size) in freshwater for 7×2013;12 mins. To deflate the swim bladder, once the operculum movement ceased, fish was removed from the freshwater and placed on a sponge. A sterile syringe was then inserted into the swim bladder from the dorsal muscle, and the plunger was slowly pulled to deflate the swim bladder (Fig. 2A).

**Fig. 2.**
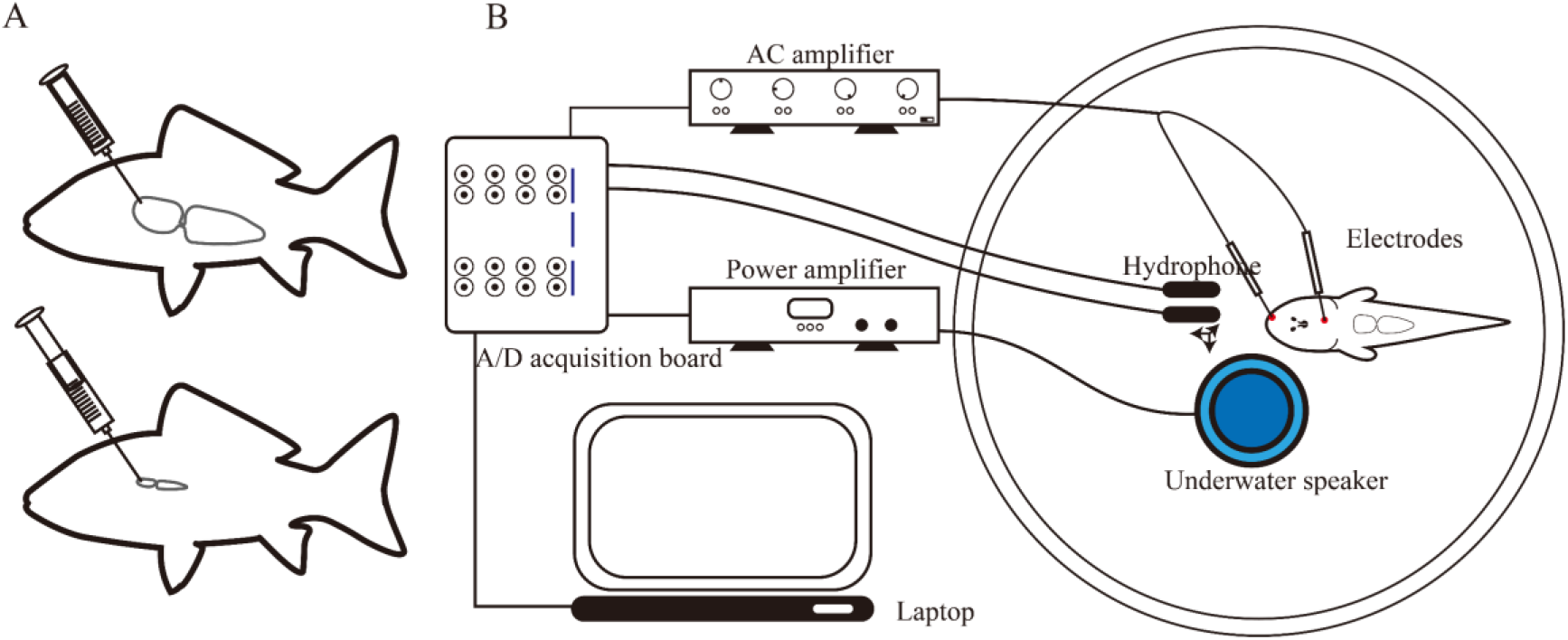
Operation for deflation of swim bladder and schematic of AEP measurement setup. (A) The swim bladder of crucian carp was deflated using a sterile syringe. (B) AEP measurement setup was composed of a laptop, an A/D acquisition board, a power amplifier, a Four-Channel Differential AC Amplifier, two hydrophones, and an underwater speaker. The experimental crucian carp was held in a customized fish holder and suspended in an acrylic tank. The AEP responses were picked up by two electroencephalography electrodes.

### AEP recording

AEP measurements were performed in an acrylic tank (0.5 m diameter, 0.3 m depth) in the Key Laboratory of Underwater Acoustic Communication and Marine Information Technology of Ministry of Education, Xiamen University, Xiamen, China. A customized fish holder consisting of a magnetic base, two plastic rods, and a clamp was used to immobilize and stabilize the fish. Muscle relaxants were not used in this study. Before experiment, each sample was anesthetized. The fish body was enveloped in a piece of medical gauze to avoid potential damage and then affixed to the clamp of the customized fish holder at a perpendicular angle. The operculum of each fish was left free to maintain normal respiration. The fish was fully submerged below the water surface. An underwater speaker (UW-30; Electro-Voice, Buchanan, MI, USA) was positioned at the center of the acrylic tank 20 cm below the fish. Three stainless-steel electrodes (single-channel electrodes, 0.8 mm diameter; KEDOUBC, China) were used to record auditory responses to sound stimuli. A recording electrode was inserted into the skull of the fish from the dorsal side. A reference electrode was inserted into the fish nares, and a ground electrode was connected to the ground using copper to avoid background interference (Fig. 2B).

The AEP measurement setup comprised a laptop (ThinkPad E430c; Lenovo, China), an A/D acquisition board (USB-6356; National Instruments, Austin, TX, USA), a power amplifier (ATA-4000; Aigtek, Xian, China), a Four-Channel Differential AC Amplifier (Model-1700; A-M Systems, Inc., Olympia, WA, USA), two hydrophones (RHSA-10; China Shipbuilding Industry Group Co., Ltd., Hangzhou, China; a flat sensitivity of −180.53 dB re 1V/μPa from 0.02 to 200 kHz), and an underwater speaker. Sound stimuli of different frequencies were generated using customized LabVIEW software programs (National Instruments, Austin, TX, USA) and then transformed into analog signals using the A/D acquisition board. This board was connected to a power amplifier/attenuator to control the output amplitude of sound stimuli before fed into the underwater speaker. The stainless-steel electrodes were connected to a four-channel differential AC amplifier, which can be set adaptively to amplify and filter the AEP signals in demand.

Modulated tone bursts were used as sound stimuli with frequency ranging from 80 to 1000 Hz (80, 100, 150, 200, 250, 300, 350, 400, 500, 600, 700, 800, 900, and 1000 Hz), covering the sensitive frequency bandwidth of the same taxa (Amoser and Ladich, 2005; Kenyon et al., 1998; Kojima et al., 2005; Vetter et al., 2018). To begin, the sound stimuli was set much higher than the ambient noise. After capturing the AEP response, sound stimuli would be attenuated at a step of 5 dB, and when it came to the proximity of the threshold, the step was changed to 2 dB. AEP waveforms recorded from a dead fish was used as a control. The stimuli were repeated 1000 times to obtain the average AEP as the final output and the stimuli phase was reversed between the first 500 and the second 500 stimuli (Stanley et al., 2020). The auditory threshold was defined as the lowest sound pressure level at which observable AEP response was detected (Kenyon et al., 1998).

### Calibration of sound stimuli

The same sound stimuli were presented for calibration after the AEP measurement. A hydrophone was fixed at the same position where the fish was placed during AEP recordings. Another hydrophone was moved to create three orthogonal axes with a fixed hydrophone to measure the accelerations in different directions. These hydrophones were connected to a laptop through the A/D acquisition board, and the data was saved using customized LabVIEW programs. Sound pressure level (SPLp-p) and sound acceleration level (SALp-p) were calibrated as:

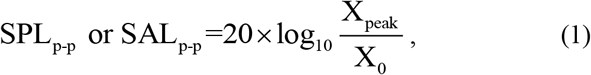

where X_peak_ represents the maximum absolute μPa or m·s^-2^ covering the given measuring period, and X_0_ = 1 μPa or m·s^-2^ is the reference value.

The acceleration was calculated using pressure data from the two hydrophones. The acceleration was determined as follows:

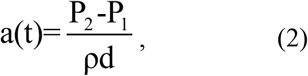

where *P* = *P_2_ - P_1_* is the sound pressure difference measured by the two hydrophones; ρ (997.8 kg m^-3^) is the density of water, and *d* is the distance between the two hydrophones. The magnitude of the total acceleration is calculated following:

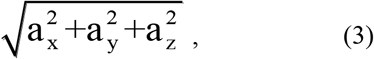

Thus, the hearing thresholds of sound pressure and acceleration can be determined.

### Numerical model development and simulation

After each AEP measurement, the fish was directly transferred to the Core Facility of Biomedical Sciences, Xiamen University for Micro-CT scanning using a SkyScan 1272 scanner (Bruker, Germany) with a resolution of 6 μm. The X-ray scanning files of different fishes were processed using the accessory software. The swim bladder and otoliths were reconstructed into individual three-dimensional (3D) models. The digital density differed among soft tissues, swim bladder (air), and solid otoliths (lapillus, sagitta, and asteriscus), which were referenced to facilitate segmentation. Stereolithography files were then generated and imported into MeshLab software (Visual Computing Lab-ISTI-CNR) to smooth the facial factor for further numerical simulations in the finite element meshes.

Based on the 3D reconstructions, we developed a numerical model for each fish and ran simulations to estimate the pressure fields in water and swim bladder, as well as the acceleration in solid otoliths. Wave propagation in an inhomogeneous fluid medium (air and water) can be computed using the following equation (Bruneau, 2013; Kinsler et al., 1999):

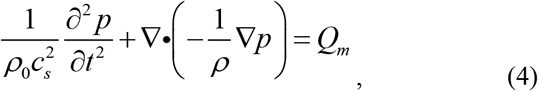

where *ρ_0_* and *C_s_* represent the static density and sound speed of the medium, respectively. *p* represents the dynamic sound pressure after the source is induced to radiate sounds, and *ρ* is the resulting variable density in the medium. To incorporate both compressional and shear wave propagations in solid otoliths, the equation was changed: (Bruneau, 2013; Rose, 1999):

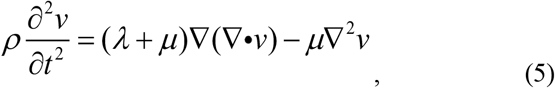

where *λ* and *μ* are two Lamé constants, and *v* is the velocity vector. The initial sound source *Q_m_* is given as a point source in the form of

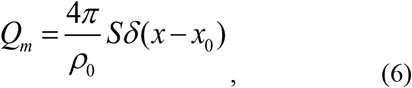

where *S* has a form of

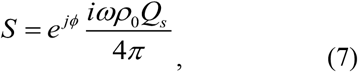

and *Q_s_* give the volume flow rate from source at *x* = *x*_0_.

To couple the interaction between the fluid and solid media, the pressure continuity condition should satisfy F*p* = **-n***p*, where F_*p*_ is the pressure load on the boundary, *p* is the pressure, and **n** is the unit normal vector. In addition, the normal acceleration *a_n_* of the acoustic pressure acting on the boundaries should be equal to the second derivative of the solid displacements. The boundary radiation was set as spherical spreading, allowing further propagation of waves with minimal reflections.

The numerical simulation was conducted using the COMSOL Multiphysics software (Stockholm, Sweden). The swim bladder and otoliths of each fish were imported into COMSOL (Fig. 3). These structures were placed in a computing sphere domain with a radius of 120 mm, creating a total volume 50 times larger than that of the swim bladder. The swim bladder was placed at the center of the computing domain. The sound speeds and densities of water and the swim bladder were set as 1481 m s^-1^ and 1000 kg m^-3^, and 343 m s^-1^ and 1.204 kg m^-3^, respectively. The otoliths were assigned a compressional wave speed of 3000 m s^-1^, a shear wave speed of 1400 m s^-1^, and a density of 2700 kg m^-3^ (Salas et al., 2019). The Pressure Acoustic-Frequency Domain module was used to solve problems in the water and swim bladder domains, whereas the otoliths were considered as solid materials, in which vibrations were solved using Solid mechanics and an Acoustic-Structure Boundary module. To minimize boundary reflection, the radiation condition was set as spherical spreading. A monopole point source, with an amplitude of 1 N/m, was set as the excitation source and placed 20 mm in front of the swim bladder. The frequency was swept from 10 to 1000 Hz with a step of 10 Hz. Volumes of the swim bladders in the control and treated groups were measured to track the coincidence between resonance frequency and volume. The distances between each pair of otoliths and swim bladder were measured as well.

**Fig. 3.**
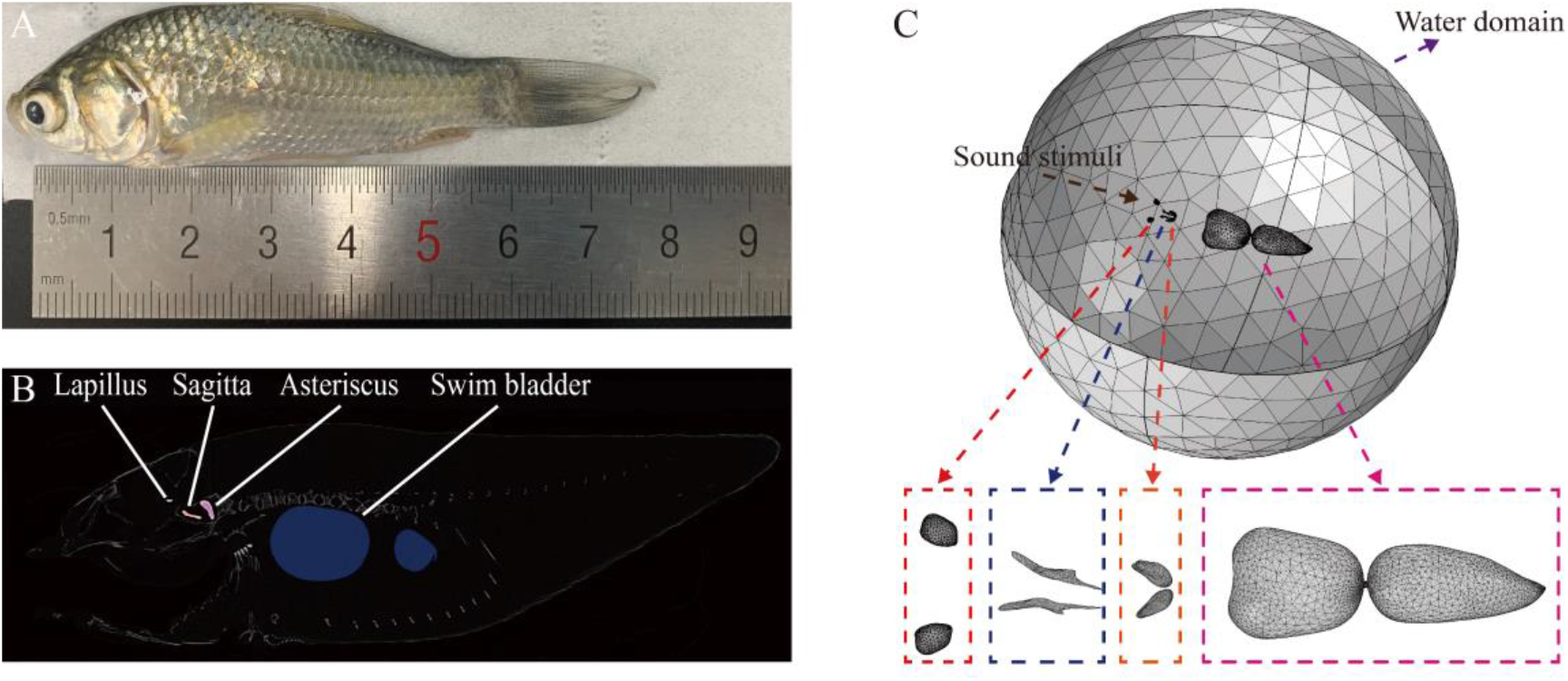
3D-reconstruction of the auditory structure in crucian carp. (A) An experimental crucian carp in vivo. (B) A sagittal cross section of crucian carp of Micro-CT imaging. (C) Setting of the numerical model includes sound stimuli, water domain, and auditory structure: lapillus (red box), sagitta (blue box), asteriscus (orange box), and swim bladder (pink box).

## RESULTS

### Influence of swim bladder on auditory threshold

The amplitude of the AEP response (Fig. 4A) decreased with the attenuation of the sound stimuli. The response from live fish was compared to that of dead fish to determine the hearing threshold at each frequency (Fig. 4B).

**Fig. 4.**
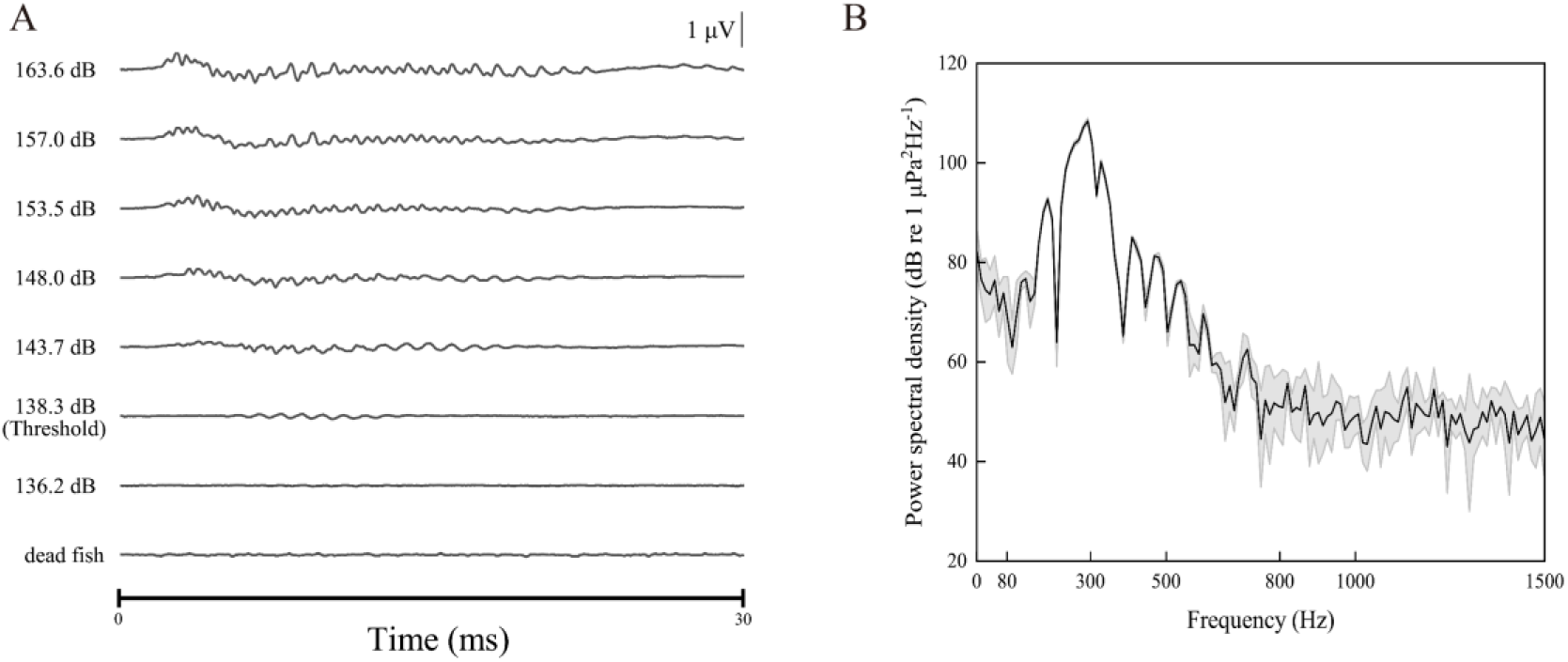
Example of AEP responses to sound stimuli at 300 Hz modulated tone bursts. (A) AEP waveforms show a definitive threshold response occurred at a sound pressure level of 138.30 dB re 1μPa. (B) Power spectral density of the sound stimuli of 300 Hz modulated tone bursts received by hydrophone.

For the five control fishes (with an intact swim bladder), the mean sound pressure threshold had a minimum of 124.18 ± 1.13 dB re 1 μPa at 100 Hz and monotonously increased to 142.36 ± 4.61 dB re 1 μPa at 600 Hz (Fig. 5A). Over this frequency, the threshold had a mean value of 138.90 ± 3.2 dB re 1 μPa, with a local lowest value at 800 Hz. At frequencies higher than 1000 Hz, no AEP response was recorded. For the five treated fishes (with a deflated swim bladder), the hearing thresholds had the lowest value (122.72 ± 1.54 dB re 1 μPa) at 80 Hz and gradually increased to the highest value (169.42 ± 0.96 dB re 1 μPa) at 800 Hz (Fig. 5A). The mean thresholds of control fishes were lower than that of treated fishes by 0.38–30.52 dB re 1 μPa (Fig. 5C). In terms of acceleration, the audiograms showed that the deflation of swim bladder led to a higher threshold, especially over 300 Hz (Figs. 5B and 5C). The AEP response was measured for all samples from 80 to1000 Hz in the control group. After deflation, the number of samples with measurable AEP response monotonously decreased from 5 below 500 Hz to 0 at 900 Hz (Fig. 5D).

**Fig. 5.**
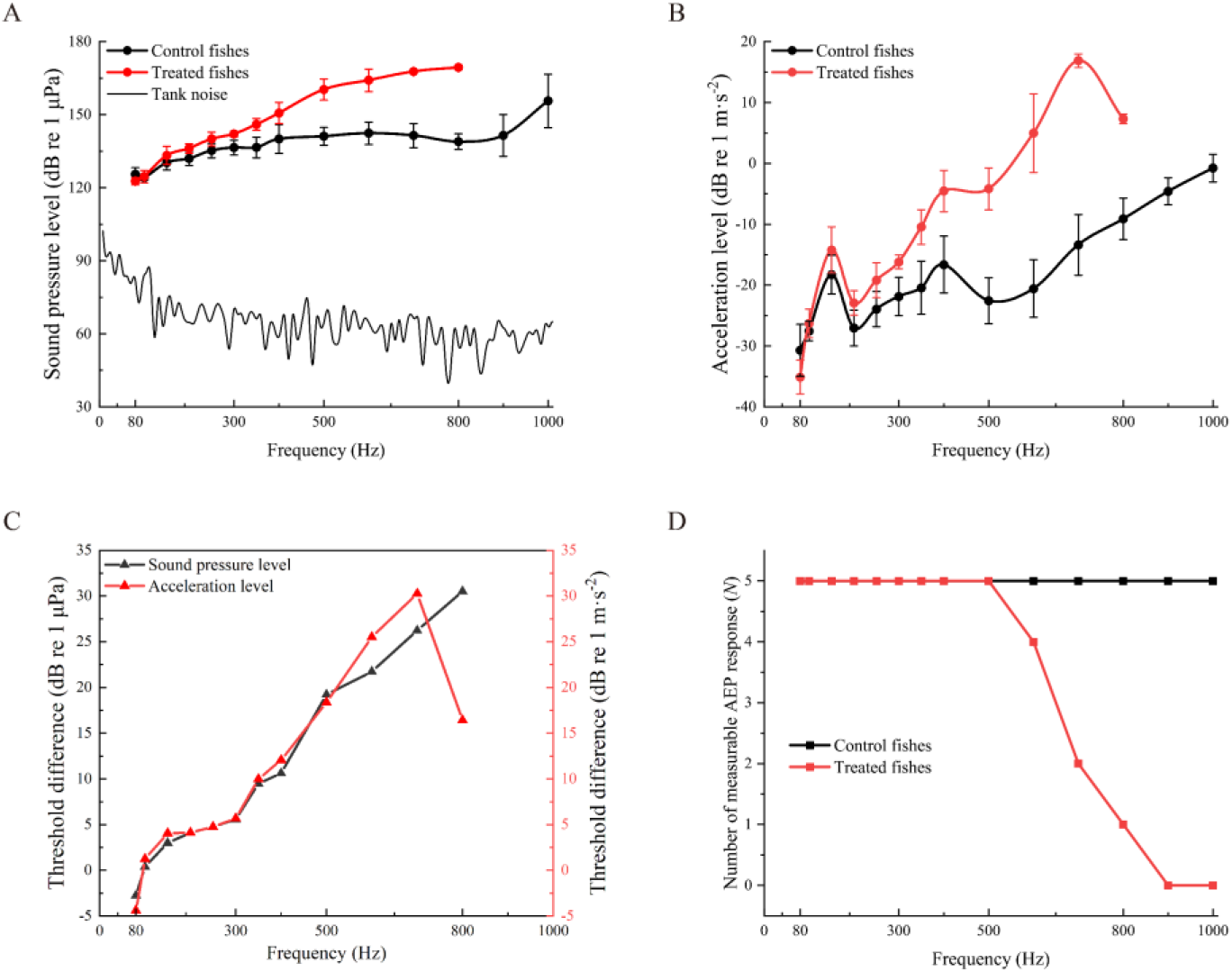
Effect of swim bladder deflation on audiograms of fishes. (A) Comparison of audiograms between intact (control fishes, *N = 5*) and deflated (treated fishes, *N = 5*) swim bladders, examined using sound pressure level and (B) comparison of audiograms using acceleration level between control and treated fishes. (C) Comparison of auditory sensitivity for control and treated fishes. (D) Number of samples with measurable AEP response at each tested frequency for control and treated fishes.

### The influence of swim bladder resonance on sound reception

The total volume of the intact swim bladder (*N = 5*) ranged from 321.74 to 359.26 mm^3^, corresponding to resonance frequencies of 840 Hz and 810 Hz, respectively. The amplitude of received sound pressure had a maximum amplitude at the resonance frequency (Fig. 6A). Volume of the deflated swim bladder (*N = 5*) ranged from 22.07 to 62.58 mm^3^ and sound pressure received in the deflated swim bladder increased monotonously within 1000 Hz (Fig. 6A). For the given example (Fig. 6A), an intact swim bladder can improve the sound pressure amplitude by 34.3 dB re 1 μPa compared with a deflated swim bladder at the resonance frequency of 840 Hz. And this difference in the mean value was 27.31 dB (*N = 5*) at 840 Hz between the control and treated groups.

**Fig. 6.**
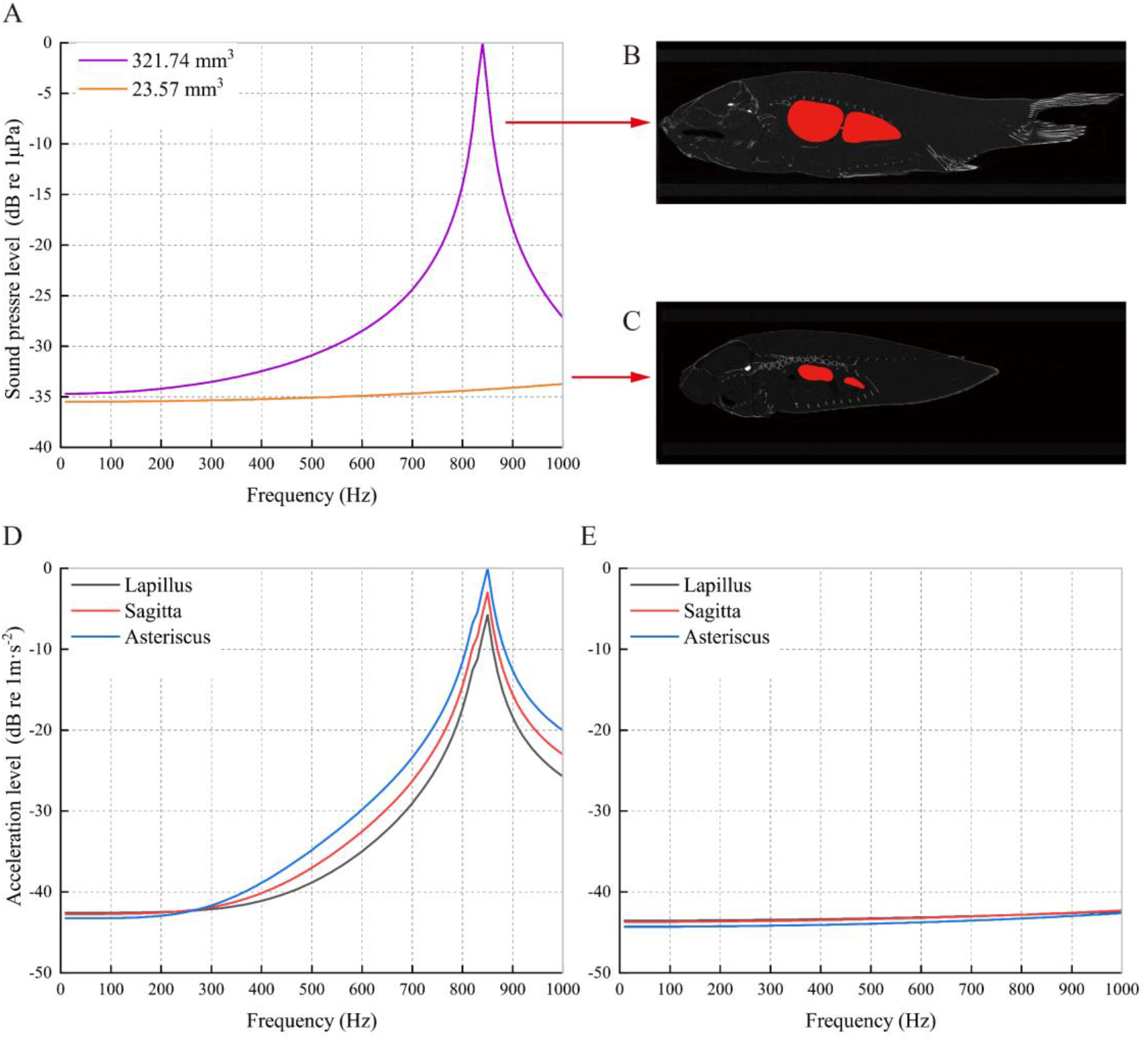
Comparison of sound reception of individuals for control fishes and treated fishes. (A) Frequency response of received sound pressure in swim bladder of an individual fish (volume, 321.74 mm^3^) in control group and a fish (volume, 23.57 mm^3^) in treated group, respectively. (B) A Micro-CT image of crucian carp with an intact swim bladder and (C) with a deflated swim bladder. (D) The acceleration of otoliths in a sample from control group and (E) treated group, respectively.

For the specimen with an intact swim bladder of 321.74 mm^3^, the acceleration was the highest in the asteriscus and second-highest in the sagittas at the resonance frequency (Fig. 6D). To compare, the vibration of otoliths in a treated fish with a deflated swim bladder of 23.57 mm^3^showed no local peaks within 1000 Hz (Fig. 6E). The acceleration of the asteriscus in the control sample was 43.13 dB re 1 m s^-2^ higher than that of the treated sample at the resonance frequency. This difference in the mean value for asteriscus was 33.78 dB (*N = 5*) between control and treated groups at resonance frequency.

## DISCUSSION

The resonance of swim bladder in crucian carp can enhance sound reception and increase auditory sensitivity, reflected in the better acoustic sensitivity and wider bandwidth in control fishes. The mean thresholds of control fishes were lower than that of treated fishes by 0.38 dB at 100 Hz and this difference was 30.52 dB at 800 Hz. This difference increased monotonously with increasing frequency. Furthermore, control fishes also had a wider bandwidth of acoustic sensitivity with a greater number of measurable AEP response over the tested frequencies of 600 to 1000 Hz. Colleye et al (2019) found that the rostral swim bladder extension in female plainfin midshipman enhanced sound pressure sensitivity and bandwidth in saccule over 305 Hz compared to fishes with a removed swim bladder (Colleye et al., 2019). A similar phenomenon was also observed in female plainfin midshipman (Vetter and Sisneros, 2020). These data suggested that low auditory threshold and wide sensitivity range were related to the presence of swim bladder.

The resonance of an intact swim bladder resulted in a local peak at the resonance frequency below 1000 Hz, which was not found for treated fishes with a deflated swim bladder. For control fishes with an intact swim bladder, the numerical simulation results were consistent with AEP recordings from 600 to 1000 Hz, both showing a local sound pressure enhancement at the resonance frequency. The auditory threshold was similar for fishes from two groups below 300 Hz (Figs. 5A, 5B, and Fig. 7), irrespective of the deflation of swim bladder, which seemingly suggested that the sensitivity was similar at low frequency. Crucian carp may have developed a diversified sound detection frequency ranges: a particle motion-sensitive region at low frequency and a sound pressure-sensitive region at high frequency.

**Fig. 7.**
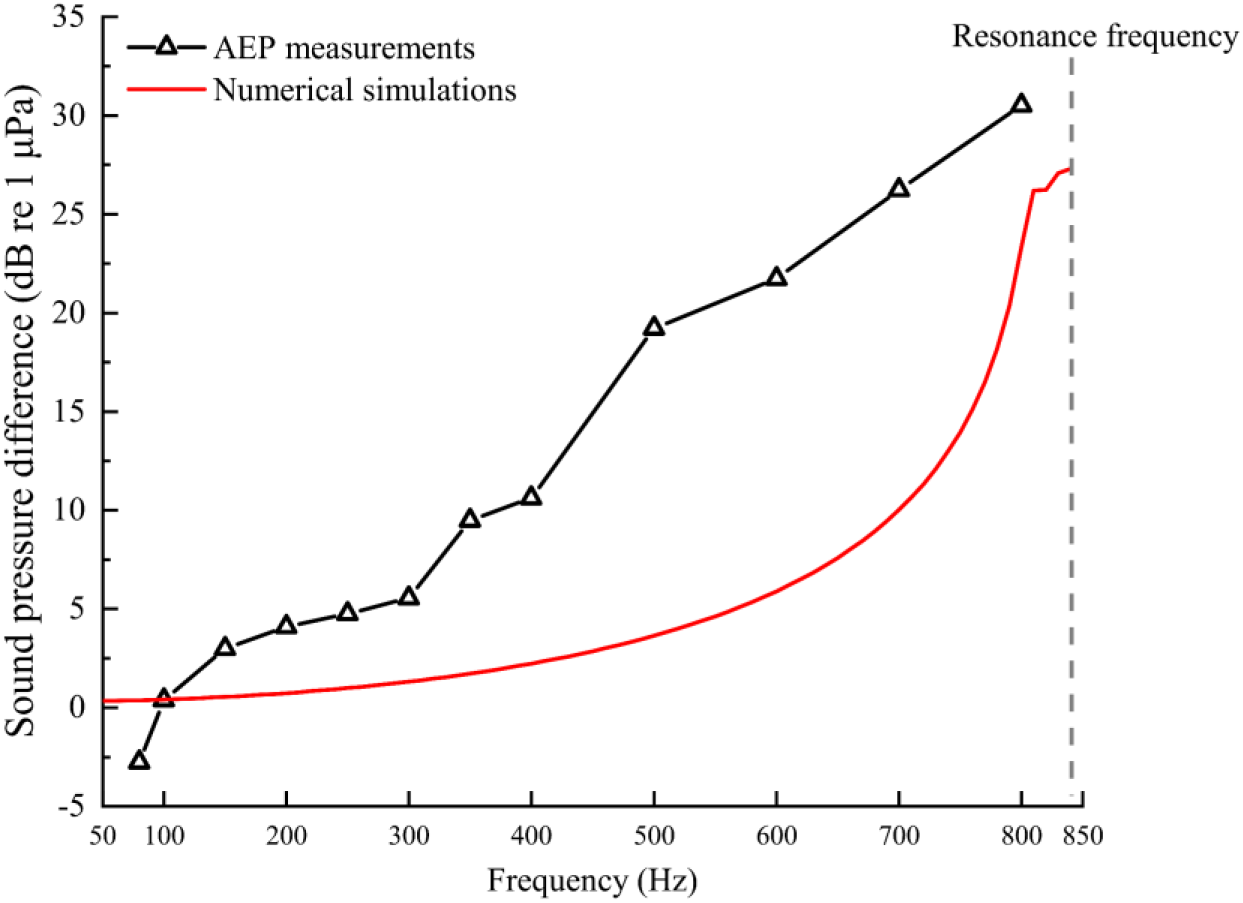
Sound pressure difference across the tested frequency range obtained from AEP measurements and numerical simulations. The data based on sound pressure level between control and treated fishes were plotted to compare the auditory sensitivity difference through AEP measurements (red) and amplitude difference obtained from numerical simulations (black).

Oscillations of the swim bladder can effectively transduce sound energy to the otoliths, especially at the resonance frequency. In teleost species, the swim bladder is thought to be a vital acoustic organ when it is sufficiently close to the otoliths or directly connects to the inner ear through acoustic coupling (Schulz-Mirbach et al., 2019). The normalized absolute sound pressure in swim bladder and the vibration in otoliths were compared between control fishes and treated fishes at 800, 820, 840 (resonance frequency), 900, and 1000 Hz (Fig. 8). The resonance of swim bladder can induce the otoliths to achieve a higher acceleration (Fig. 8A). In comparison, the otoliths had no significant vibration within the tested frequency range after deflation (Fig. 8B). Though we simplified the fish model by comprising only a swim bladder and three pairs of otoliths, the function of bladder in sound reception and hearing can be glimpsed, consistent with a previous study (Salas et al., 2019). Structures including muscles and skeletons should be incorporated in the future as well to better quantify the contributions of the swim bladder and other structures to fish hearing.

**Fig. 8.**
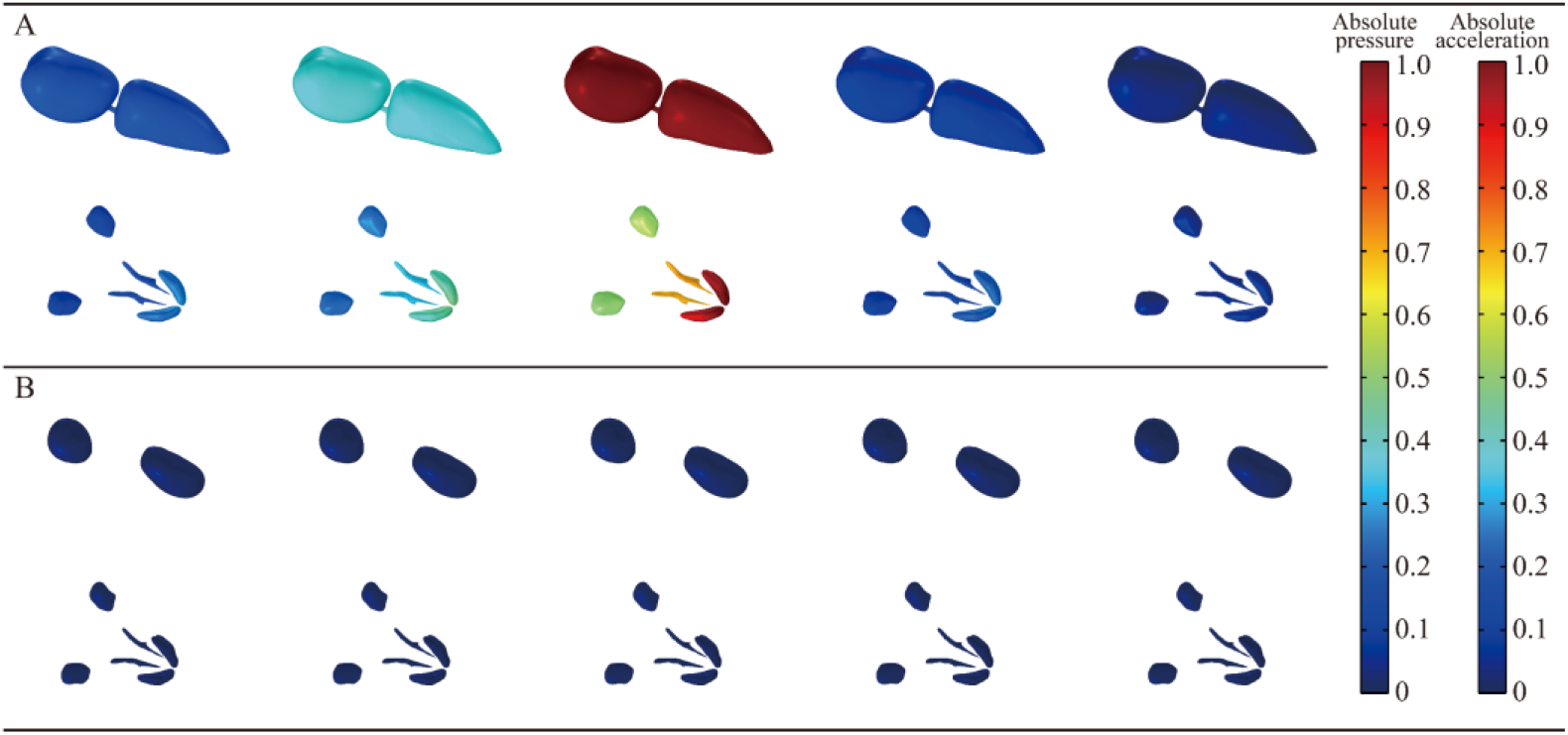
Normalized absolute sound pressure in the swim bladder and vibration in otoliths at 800, 820, 840 (resonance frequency), 900, and 1000 Hz. (A) Sound pressure field in the swim bladder and the induced-vibration of different otoliths for a control fish with an intact swim bladder (volume, 321.74 mm^3^) and (B) a treated fish with a deflated swim bladder (volume, 23.57 mm^3^).

There was an opposite relationship between resonance frequency and volume of swim bladder (Fig. 9A), which was previously addressed in red drum (*Sciaenops ocellatus*) (Salas *et al.* 2019) The swim bladder of the red drum had a volume ranging from 0.26 to 2.7 mm^3^, corresponding to the resonance frequency of 8750 and 4250 Hz (Salas et al., 2019). Here, the additional numerical simulations of the deflated swim bladder were conducted up to 5000 Hz with a step of 10 Hz. After deflation, the swim bladder was split into two individual bladders with volumes ranging from 6.59 to 40.47 mm^3^, with a low end of resonance frequency of 1580 Hz and high end of 2920 Hz. However, no AEP response was recorded from 1580 Hz and 2920 Hz in our specimens, suggesting a limited function of the swim bladder through resonance at frequencies over the hearing range or this may root in response speed of nervous system and modulation of auditory structures. For instance, the utricular specialization of the sensory epithelium (or macula) in clupeid fish is divided into three distinct parts and this specialization improves the hearing sensitivity in a number of clupeid fishes to ultrasound stimuli up to at least 180 kHz (Mann and Lobel, 1997; Mann et al., 1998; Popper et al., 2004). Future studies should locate specimens sensitive to sounds in high frequency range and incorporate the resonance and the related AEP responses to better quantify their connections.

**Fig. 9.**
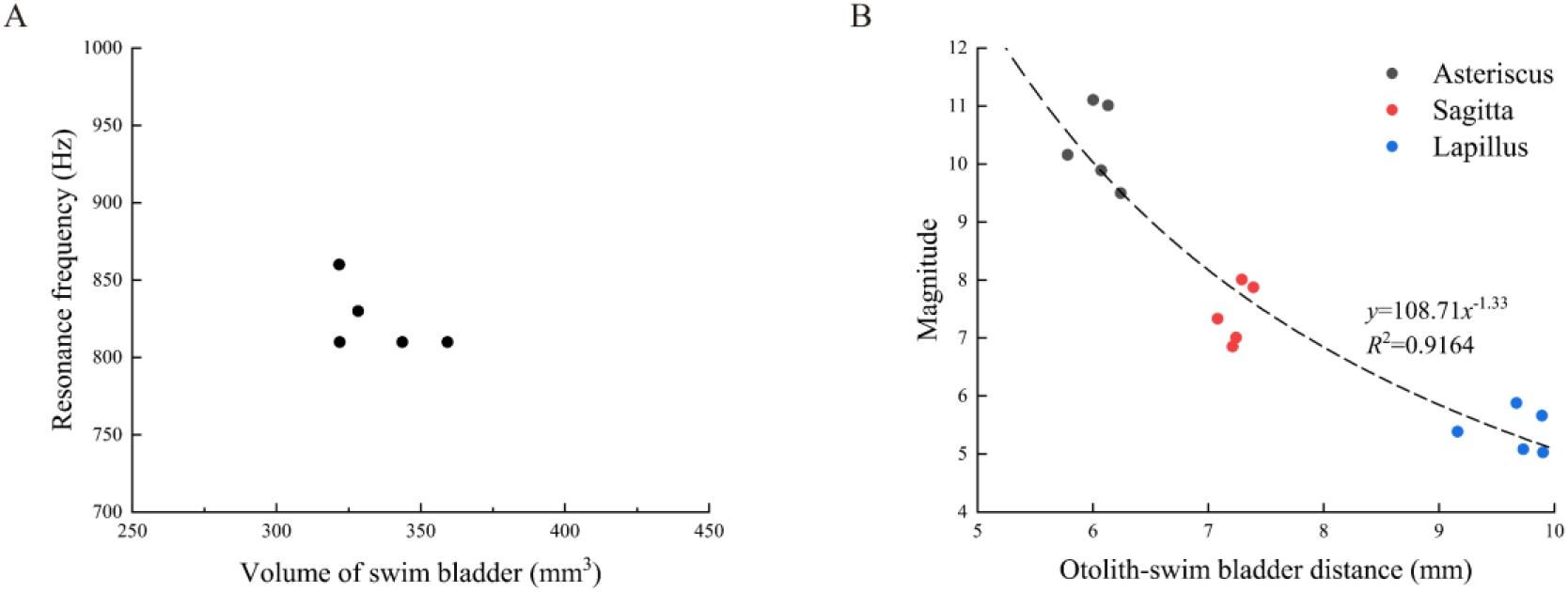
Change in crucian carp swim bladder and otoliths acoustic properties. (A) The relation between resonance frequency and swim bladder volume. (B) The relationship between otolith-swim bladder distance and acceleration magnitude in three pairs of otoliths.

The asteriscus has the maximum acceleration, followed by the sagitta and lapillus. After deflation, no local peak was found with the tested frequency range for all otoliths. At the resonance frequency, the otoliths oscillate stronger because of transduction from the swim bladder. This mechanism might indicate a difference in hearing pressure sensitivity when the swim bladder undergoes morphological changes (Schulz-Mirbach et al., 2012). The acceleration of the otoliths depended on the distance between the swim bladder and the otoliths (Fig. 9B). The asteriscus, which was closest to swim bladder distance, exhibited the highest acceleration among all otoliths. This was different from the findings in sleeper goby (*Dormitator latifrons*) and plainfin midshipman, which showed that the thresholds of the lagenar potentials were higher than those of the saccule (Lu et al., 2003; Lu et al., 2010; Vetter et al., 2019). The difference between lagenar and saccular vibration might be due to the distance from the swim bladder to the otoliths (Fig. 9B).

In our study, treated fishes had a similar sensitivity to control fishes below 300 Hz. Previous studies have indicated that the otolith end organs are sensitive to particle motion below 400 Hz (Lu et al., 2003; Lu et al., 2004; Vetter, 2019; Vetter et al., 2019). Our results may suggest that fish separate their sensing ranges into low-frequency particle motion (as a vector) and relatively high-frequency sound pressure (scalar) sensing.

Using traditional surgical methods to investigate the motion of the otoliths are challenging because of the limitation of inserting recorder into the specimen. Recently, a new experimental setup was used to visualize the motion of the isolated otoliths in 4D, providing insights into the motion of auditory structures (Maiditsch et al., 2022). Besides, finite element modeling offers an alternative way to study the roles of the otoliths in fish hearing and has been widely used in odontocetes to investigate sound transmission and reception (Cranford and Krysl, 2015; Song et al., 2021; Tubelli et al., 2018), which may facilitate more studies on hearing mechanism in fish.

We combined AEP measurement, Micro-CT scanning, and numerical simulation to show that the resonance of swim bladder can enhance pressure sensing by 34.3 dB re 1 μPa and acceleration sensing by 43.13 dB re 1 m s^-2^ at resonance frequency. Fish might have developed various means to sense particle motion and pressure at separate frequency ranges. These results help us to better understand fish hearing mechanism.

## LIST OF SYMBOLS AND ABBREVIATIONS

AEP: Auditory evoked potentials
SPLp-p: Threshold of sound pressure
SAL_p-p_: Threshold of acceleration
Micro-CT: Micro-computed tomography
3D: Three-dimensional

## ACKNOWLEDGEMENTS

We would like to acknowledge Xuguang Zhang from Shanghai Ocean University, for his professional advice in AEP experiment design and thanks are given to Chi Zhang form Ocean University of China for his assistance in identifying otoliths. Special thanks are given to all members of the MBAT Lab in the College of Ocean and Earth Sciences at Xiamen University.

## COMPETING INTERESTS

The authors declare no competing or financial interests.

## AUTHOR CONTRIBUTIONS

Conceptualization: Z. S., Y. Z.; Methodology: H. L., Z. G., Z. S., Y. S., W. O., Y. Z.; Investigation: H. L., Z. S., Y. S., J. Z.; Resources: Z. S., Y. Z.; Writing - original draft: H. L., Z. S., Z. G., Y. S.; Writing - review & editing: H. L., Z. S., Y. Z.; Project administration: Z. S., Y. Z.; Funding acquisition: Z. S., Y. Z.

## FUNDING

This work was funded by the Development of a novel high-performance platform for deep-sea fish farming and research of its key performances for Science and Technology Major Project of Fujian Province (Grant Nos. 2021NZ033016), the National Natural Science Foundation of China (Grant Nos: 12074323; 42106181), the China National Postdoctoral Program for Innovative Talents (Grant No: BX2021168), the China Postdoctoral Science Foundation (Grant No: 2020M682086), and the Outstanding Postdoctoral Scholarship, State Key Laboratory of Marine Environmental Science at Xiamen University.

## DATA AVAILABILITY

All data needed to reproduce the results in this paper are available from the corresponding authors upon reasonable request.

## Notes

### Competing Interest Statement

The authors have declared no competing interest.

## REFERENCES

Amoser, S. and Ladich, F. (2005). Are hearing sensitivities of freshwater fish adapted to the ambient noise in their habitats? J. Exp. Biol. 208, 3533–3542.

Blaxter, J. H. S. and Fuiman, L. A. (1990). The role of the sensory systems of herring larvae in evading predatory fishes. J. Mar. Biol. Assoc. U. K. 70, 413–427.

Bruneau, M. (2013). Fundamentals of Acoustics (Wiley, New York).

Colleye, O., Vetter, B. J., Mohr, R. A., Seeley, L. H. and Sisneros, J. A. (2019). Sexually dimorphic swim bladder extensions enhance the auditory sensitivity of female plainfin midshipman fish, Porichthys notatus. J. Exp. Biol. jeb.204552.

Corwin, J. T., Bullock, T. H. and Schweitzer, J. (1982). The auditory brain stem response in five vertebrate classes. Electroencephalogr. Clin. Neurophysiol. 54, 629–641.

Cranford, T. W. and Krysl, P. (2015). Fin Whale Sound Reception Mechanisms: Skull Vibration Enables Low-Frequency Hearing. PLOS ONE 10, e0116222.

Fay, R. R. (1984). The Goldfish Ear Codes the Axis of Acoustic Particle Motion in Three Dimensions. Science 225, 951–954.

Giorli, G., Au, W. W. L. and Neuheimer, A. (2016). Differences in foraging activity of deep sea diving odontocetes in the Ligurian Sea as determined by passive acoustic recorders. Deep Sea Res. Part Oceanogr. Res. Pap. 107, 1–8.

Gordon, T. A. C., Harding, H. R., Wong, K. E., Merchant, N. D., Meekan, M. G., McCormick, M. I., Radford, A. N. and Simpson, S. D. (2018). Habitat degradation negatively affects auditory settlement behavior of coral reef fishes. Proc. Natl. Acad. Sci. 115, 5193–5198.

Hawkins, A. D. and Amorim, M. C. P. (2000). Spawning Sounds of the Male Haddock, Melanogrammus aeglefinus. Environ. Biol. Fishes 59, 29–41.

Horodysky, A. Z., Brill, R. W., Fine, M. L., Musick, J. A. and Latour, R. J. (2008). Acoustic pressure and particle motion thresholds in six sciaenid fishes. J. Exp. Biol. 211, 1504–1511.

Hudspeth, A. J. (1985). The Cellular Basis of Hearing: The Biophysics of Hair Cells. Science 230, 745–752.

Kenyon, T. N., Ladich, F. and Yan, H. Y. (1998). A comparative study of hearing ability in fishes: the auditory brainstem response approach. J. Comp. Physiol. [A] 182, 307–318.

Kinsler, P., Kelsall, R. W. and Harrison, P. (1999). Interface and confined phonons in stepped quantum wells. Phys. B Condens. Matter 263–264, 507–509.

Kojima, T., Ito, H., Komada, T., Taniuchi, T. and Akamatsu, T. (2005). Measurements of auditory sensitivity in common carp *Cyprinus carpio* by the auditory brainstem response technique and cardiac conditioning method. Fish. Sci. 71, 95–100.

Krause, B. B. (2008). Anatomy of the Soundscape Evolving Perspectives. J. Audio Eng. Soc. Audio Eng. Soc. 56, 73–80.

Ladich, F. (2016). Peripheral Hearing Structures in Fishes: Diversity and Sensitivity of Catfishes and Cichlids. In Fish Hearing and Bioacoustics (ed. Sisneros, J. A.), pp. 321–340. Cham: Springer International Publishing.

Ladich, F. and Fay, R. R. (2013). Auditory evoked potential audiometry in fish. Rev. Fish Biol. Fish. 23, 317–364.

Lu, Z., Xu, Z. and Buchser, W. J. (2003). Acoustic response properties of lagenar nerve fibers in the sleeper goby, Dormitator latifrons. J. Comp. Physiol. [A] 189, 889–905.

Lu, Z., Xu, Z. and Buchser, W. J. (2004). Coding of acoustic particle motion by utricular fibers in the sleeper goby, Dormitator latifrons. J. Comp. Physiol. [A].

Lu, Z., Xu, Z. and Buchser, W. J. (2010). Frequency coding of particle motion by saccular afferents of a teleost fish. J. Exp. Biol. 213, 1591–1601.

Maiditsch, I. P. and Ladich, F. (2022). Acoustic and visual adaptations to predation risk: a predator affects communication in vocal female fish. Curr. Zool. 68, 149–157.

Maiditsch, I. P., Ladich, F., Heß, M., Schlepütz, C. M. and Schulz-Mirbach, T. (2022). Revealing sound-induced motion patterns in fish hearing structures in 4D: a standing wave tube-like setup designed for high-resolution time-resolved tomography. J. Exp. Biol. 225, jeb243614.

Mann, D. A. and Lobel, P. S. (1997). Propagation of damselfish *(Pomacentridae)* courtship sounds. J. Acoust. Soc. Am. 101, 3783–3791.

Mann, D. A., Lu, Z., Hastings, M. C. and Popper, A. N. (1998). Detection of ultrasonic tones and simulated dolphin echolocation clicks by a teleost fish, the American shad (Alosa sapidissima). J. Acoust. Soc. Am. 104, 562–568.

Miller, L. A., Simon, M., Ugarte, F. and Wahlberg, M. (2006). Exploitation of sound during predator - prey interactions: Killer whales and herring. J. Acoust. Soc. Am. 119, 3372–3372.

Montgomery, J., Jeffs, A., Simpson, S., Meekan, M. and Tindle, C. (2006). Sound as an orientation clue for the pelagic larvae of reef fish and crustaceans. Adv. Mar. Biol. 51, 143–196.

Mooney, T. A., Hanlon, R. T., Christensen-Dalsgaard, J., Madsen, P. T., Ketten, D. R. and Nachtigall, P. E. (2010). Sound detection by the longfin squid *(Loligo pealeii)* studied with auditory evoked potentials: sensitivity to low-frequency particle motion and not pressure. J. Exp. Biol. 213, 3748–3759.

Parmentier, E., Berten, L., Rigo, P., Aubrun, F., Nedelec, S. L., Simpson, S. D. and Lecchini, D. (2015). The influence of various reef sounds on coral-fish larvae behaviour: reef-sound influence on fish larvae behaviour. J. Fish Biol. 86, 1507–1518.

Popper, A. N. and Fay, R. R. (1993). Sound Detection and Processing by Fish: Critical Review and Major Research Questions (Part 2 of 2). Brain. Behav. Evol. 41, 26–38.

Popper, N. and Fay, R. (1997). Evolution of the Ear and Hearing: Issues and Questions. Brain. Behav. Evol. 50, 213–221.

Popper, A. N. and Fay, R. R. (2011). Rethinking sound detection by fishes. Hear. Res. 273, 25–36.

Popper, A. N., Fay, R. R., Platt, C. and Sand, O. (2003). Sound Detection Mechanisms and Capabilities of Teleost Fishes. In Sensory Processing in Aquatic Environments (ed. Collin, S. P. and Marshall, N. J.), pp. 3–38. New York, NY: Springer New York.

Popper, A. N., Plachta, D. T. T., Mann, D. A. and Higgs, D. (2004). Response of clupeid fish to ultrasound: a review. ICES J. Mar. Sci. 61, 1057–1061.

Rogers, P. H. and Cox, M. (1988). Underwater Sound as a Biological Stimulus. In Sensory Biology of Aquatic Animals (ed. Atema, J., Fay, R. R., Popper, A. N., and Tavolga, W. N.), pp. 131–149. New York, NY: Springer New York.

Rose, J. L. (1999). Ultrasonic waves in solid media. Cambridge; New York: Cambridge University Press.

Salas, A. K., Wilson, P. S. and Fuiman, L. A. (2019). Ontogenetic change in predicted acoustic pressure sensitivity in larval red drum (Sciaenops ocellatus). J. Exp. Biol. jeb.201962.

Schulz-Mirbach, T., Metscher, B. and Ladich, F. (2012). Relationship between Swim Bladder Morphology and Hearing Abilities-A Case Study on Asian and African Cichlids. PLoS ONE 7, e42292.

Schulz-Mirbach, T., Ladich, F., Plath, M. and Heß, M. (2019). Enigmatic ear stones: what we know about the functional role and evolution of fish otoliths: The role of fish otoliths in inner ear function. Biol. Rev. 94, 457–482.

Shao, Y. T., Chen, I.-S. and Yan, H. Y. (2014). The auditory roles of the gas bladder and suprabranchial chamber in walking catfish (Clarias batrachus). Zool. Stud. 53, 1.

Sisneros, J. A. and Rogers, P. H. (2016). Directional Hearing and Sound Source Localization in Fishes. In Fish Hearing and Bioacoustics (ed. Sisneros, J. A.), pp. 121–155. Cham: Springer International Publishing.

Song, Z., Zhang, J., Ou, W., Zhang, C., Dong, L., Dong, J., Li, S. and Zhang, Y. (2021). Numerical-modeling-based investigation of sound transmission and reception in the short-finned pilot whale (*Globicephala macrorhynchus*). J. Acoust. Soc. Am. 150, 225–232.

Stanley, J. A., Caiger, P. E., Phelan, B., Shelledy, K., Mooney, T. A. and Van Parijs, S. M. (2020). Ontogenetic variation in the hearing sensitivity of black sea bass *(Centropristis striata)* and the implications of anthropogenic sound on behavior and communication. J. Exp. Biol. jeb.219683.

Sueur, J. and Farina, A. (2015). Ecoacoustics: the Ecological Investigation and Interpretation of Environmental Sound. Biosemiotics 8, 493–502.

Thode, A. M., Sakai, T., Michalec, J., Rankin, S., Soldevilla, M. S., Martin, B. and Kim, K. H. (2019). Displaying bioacoustic directional information from sonobuoys using “azigrams.” J. Acoust. Soc. Am. 146, 95–102.

Tubelli, A. A., Zosuls, A., Ketten, D. R. and Mountain, D. C. (2018). A model and experimental approach to the middle ear transfer function related to hearing in the humpback whale (Megaptera novaeangliae). J. Acoust. Soc. Am. 144, 525–535.

Van Oosterom, L., Montgomery, J. C., Jeffs, A. G. and Radford, C. A. (2016). Evidence for contact calls in fish: conspecific vocalisations and ambient soundscape influence group cohesion in a nocturnal species. Sci. Rep. 6, 19098.

Vetter, B. J. (2019). Role of the Lagena in fish hearing and its susceptibility to anthropogenic noise. p. 010001. Den Haag, The Netherlands.

Vetter, B. J. and Sisneros, J. A. (2020). Swim bladder enhances lagenar sensitivity to sound pressure and higher frequencies in female plainfin midshipman (*Porichthys notatus*). J. Exp. Biol. jeb.225177.

Vetter, B. J., Brey, M. K. and Mensinger, A. F. (2018). Reexamining the frequency range of hearing in silver *(Hypophthalmichthys molitrix)* and bighead (*H. nobilis)* carp. PLOS ONE 13, e0192561.

Vetter, B. J., Seeley, L. H. and Sisneros, J. A. (2019). Lagenar potentials of the vocal plainfin midshipman fish, Porichthys notatus. J. Comp. Physiol. [A] 205, 163–175.

Yan, H. Y., Fine, M. L., Horn, N. S. and Colon, W. E. (2000). Variability in the role of the gasbladder in fish audition. J. Comp. Physiol. [A] 186, 435–445.

